# Microbial Diversity of Deep-Sea Ferromanganese Crust Field in the Rio Grande Rise, Southwestern Atlantic Ocean

**DOI:** 10.1101/2020.06.13.150011

**Authors:** Natascha Menezes Bergo, Amanda Gonçalves Bendia, Juliana Correa Neiva Ferreira, Bramley Murton, Frederico Pereira Brandini, Vivian Helena Pellizari

**Affiliations:** Department of Biological Oceanography, Oceanograophic Institute, University of Sao Paulo, Brazil; Natural Environment Research Council, England

**Author notes:** **Corresponding author** Correspondence to Dra Vivian Helena Pellizari. **Availability of data and material** The sequencing reads generated for this study can be found in the National Centre for Biotechnology Information (NCBI) database under the BioProject PRJNA638744.

**Keywords:** Deep-sea Ferromanganese Crusts, Microbial Community, Biogeochemical Cycling, Rio Grande Rise, Geomicrobiology

## Abstract

Seamounts are often covered with Fe and Mn oxides, known as ferromanganese (Fe-Mn) crusts. Future mining of these crusts is predicted to have significant effects on biodiversity in mined areas. Although microorganisms have been reported on Fe–Mn crusts, little is known about the role of crusts in shaping microbial communities. Here, we investigated microbial community based on 16S rRNA gene sequences retrieved from Fe-Mn crusts, coral skeleton, calcarenite and biofilm at crusts of the Rio Grande Rise (RGR). RGR is a prominent topographic feature in the deep southwestern Atlantic Ocean with Fe-Mn crusts. Our results revealed that crust field of the RGR harbors a usual deep-sea microbiome. We observed differences of microbial structure according to the sampling location and depth, suggesting an influence of water circulation and availability of particulate organic matter. Bacterial and archaeal groups related to oxidation of nitrogen compounds, such as Nitrospirae, Nitrospinae phyla, Nitrosopumilus within Thaumarchaeota group were present on those substrates. Additionally, we detected abundant assemblages belonging to methane oxidation, i. e. Ca. Methylomirabilales (NC10) and SAR324 (Deltaproteobacteria). The chemolithoautotrophs associated with ammonia-oxidizing archaea and nitrite-oxidizing bacteria potentially play an important role as primary producers in the Fe-Mn substrates from RGR. These results provide the first insights into the microbial diversity and potential ecological processes in Fe-Mn substrates from the Atlantic Ocean. This may also support draft regulations for deep-sea mining in the region.

## INTRODUCTION

The process of biogenesis in Fe-Mn crusts formation is not well described at all. Microorganisms are suggested to be potentially involved in the elemental transition between seawater and Fe-Mn substrates [15, 43], they might play a role in Mn oxidation and precipitation. Microbial diversity and abundance have been studied in the Fe-Mn crusts, and their associated sediment and water of Pacific Ocean seamounts [16, 24, 30,31]. These have detected the potential for microbial chemosynthetic primary production supported by ammonia oxidation [12, 15, 31]. Wang and Muller (2009) proposed that free-living and biofilm-forming bacteria provide the matrix for manganese deposition, and coccolithophores are the dominant organisms that act as bio-seeds for an initial manganese deposition in the Fe-Mn crusts. Kato et al, (2019) proposed a model of the microbial influence in the biogeochemical cycling of C, N, S, Fe and Mn in Fe-Mn crusts, which indicates their contribution to crust development. In contrast, other authors suggested that these endolithic and epilithic microbial communities just live in/on Fe-Mn crust as a favorable physical substrate [12, 30].

Fe-Mn crust occurs on rocky substrates at seamounts globally and has a slow accretion rate of typically 1–5 mm per million years [10]. Seamounts and rises are most common in the Pacific Ocean where there are estimated to be over 50,000 [19, 44]. Fe-Mn crusts are economically important because they mainly contain cobalt and other rare and trace metals of high demand including copper, nickel, platinum, and tellurium [10]. While crusts are potentially of economic value, seamounts also offer a habitat appropriate for sessile fauna, most corals and sponges [9, 35]. However, the ecological roles of chemosynthetic and heterotrophic microbes in metal-rich benthic ecosystems remain unknown for the Atlantic Ocean.

Despite the increasing efforts that have been made using high-throughput DNA sequencing of microbes living on Fe-Mn crusts from the Pacific Ocean, similar studies from the Atlantic Ocean are still scarce. Considering the wide distribution of Fe-Mn crusts on seamounts globally and their potential for future mineral resource supplies [10], the study of their microbiome, function and resilience to environmental impacts caused by its exploitation is essential [33].

To better understand how microbial community structure is shaped by metallic substrates in seamounts, we sampled Fe-Mn crust biofilm, crusts, encrusted coral skeletons, and calcarenite substrate from the Rio Grande Rise (RGR), Southwestern Atlantic Ocean. We used the sequencing of the 16S rRNA gene to determine the microbial diversity and community structure, and to predict microbial functions and ecological processes. We hypothesized that there is a core microbiome among the Fe-Mn substrates, despite the influence of different environmental conditions such as water masses, temperature, salinity and depth. Our results revealed a diverse microbial community among the Fe-Mn substrates, including groups assigned within Thaumarchaeota, *Gammaproteobacteria* and *Alphaproteobacteria*. Although differences of microbial diversity were noted according to the sampling location and depth, we confirmed the presence of a core microbiome among the Fe-Mn substrates mainly composed by ammonia-oxidizing Archaea.

## MATERIAL AND METHODS

### Field Sampling

The study area, RGR, is an extensive topographic feature of ~150,000km^2^ in the Southwestern Atlantic Ocean [7] (Fig. 1). RGR is approximately 1,000 – 4,000m deep and is located ~1,000km to the east of the Brazil and Argentine basins [7, 28].

**Fig. 1.**
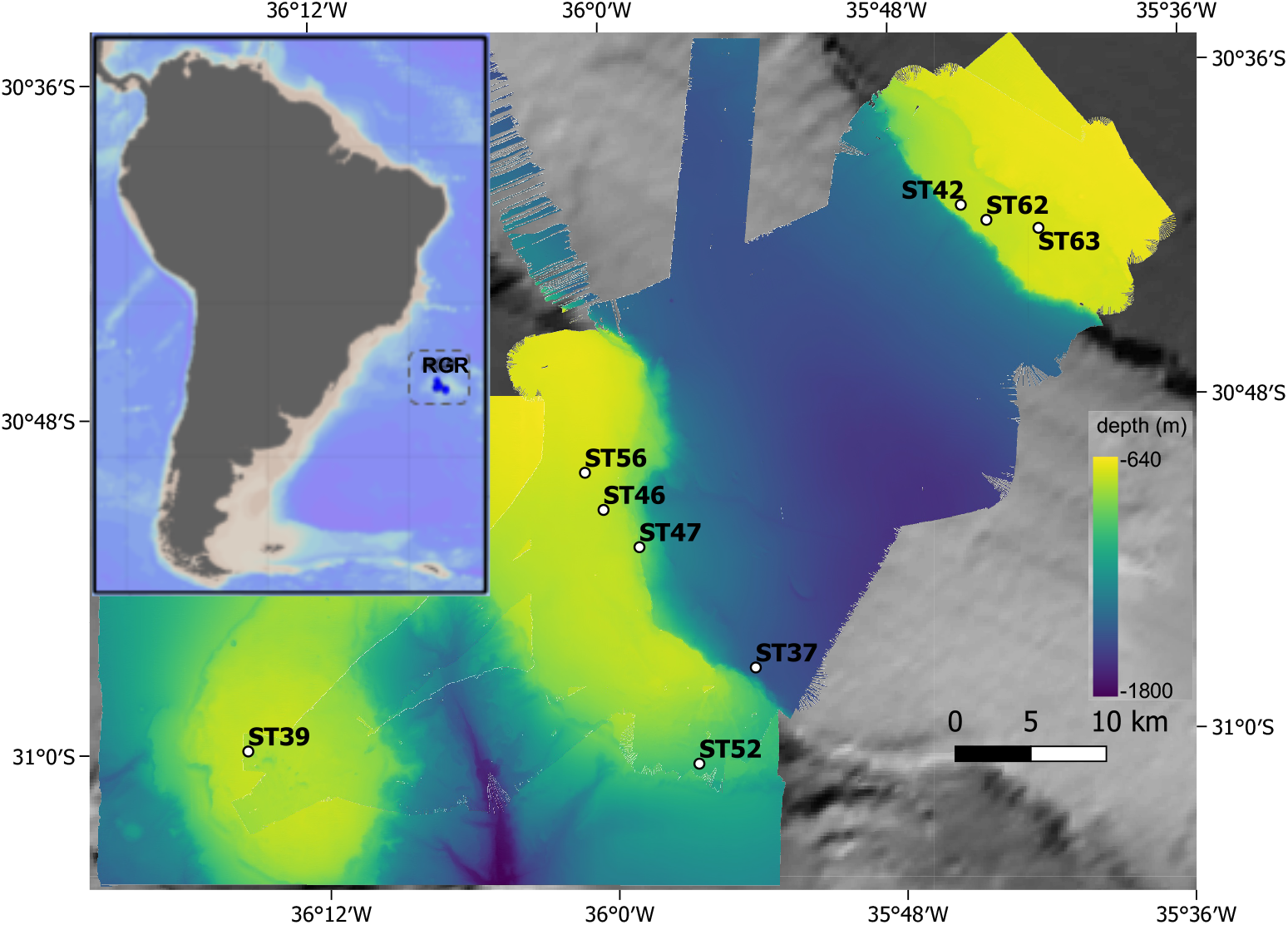
Study area and collection stations during the oceanographic expedition RGR2 to the RGR, Atlantic Ocean. High resolution bathymetry performed during the cruise

Fe-Mn crust biofilm, crust, encrusted coral skeletons, and calcarenite samples were collected during the Marine E-Tech RGR2 expedition DY094 onboard the Royal Research Ship (RRS) Discovery (NOC-NERC, UK) in October 2018 from the RGR (Supplementary Figure 1 and Table 1). Once onboard the vessel, the recovered material was aseptically sampled from each dredge. The surface of the crust and encrusted corals samples was washed with seawater filtered through a 0.2μm-pore polycarbonate membrane to remove loosely attached particles, and immediately stored in sterile Whirl-Pak bags at −80°C.

**Table 1.** Samples list, sample type, latitude, longitude and depth from RGR.

| Sample ID | Sample Type | Water mass | Station | LAT | LONG | Depth (m) |
| --- | --- | --- | --- | --- | --- | --- |
| DY094_63_01 | Fe-Mn Crust | SACW | 63 | 30.704 | 35.704 | 663 |
| DY094_56_01 | Fe-Mn Crust | SACW | 56 | 30.838 | 36.016 | 664 |
| DY094_47_01 | Fe-Mn Crust | SACW | 47 | 30.875 | 35.981 | 762 |
| DY094_46_01 | Fe-Mn Crust | SACW | 46 | 30.854 | 35.005 | 685 |
| DY094_47_02 | Fe-Mn encrusted coral skeletons | SACW | 47 | 30.875 | 35.981 | 762 |
| DY094_52_02 | Fe-Mn encrusted coral skeletons | AIA | 52 | 31.009 | 35.943 | 903 |
| DY094_62_01 | Calcarenite | SACW | 62 | 30.698 | 35.742 | 661 |
| DY094_62_02 | Calcarenite | SACW | 62 | 30.698 | 35.742 | 661 |
| DY094_39_01 | Fe-Mn Crust Biofilm | SACW | 37 | 30.969 | 35.912 | 881 |
| DY094_45_01 | Fe-Mn Crust Biofilm | SACW | 39 | 30.005 | 35.245 | 758 |
| DY094_46_03 | Fe-Mn Crust Biofilm | SACW | 46 | 30.854 | 35.005 | 685 |
| DY094_61_01 | Fe-Mn Crust Biofilm | SACW | 42 | 30.689 | 35.746 | 711 |

### DNA Extraction, 16S rRNA gene Amplification and Sequencing

DNA was extracted from the recovered samples using 5g of samples with the FastDNA™ SPIN Kit for Soil (MPBiomedical), according to the manufacturer’s protocol [21, 38]. DNA integrity was determined after electrophoresis in 1% (v/v) agarose gel prepared with TAE 1X (Tris 0.04M, glacial acetic acid 1M, EDTA 50mM, pH 8), and staining with Sybr Green (Thermo Fisher Scientific, São Paulo, Brazil). DNA concentration was determined using the Qubit dsDNA HS assay kit (Thermo Fisher Scientific, São Paulo, Brazil), according to the manufacturer’s instructions.

Before sending samples for preparation of Illumina libraries and sequencing, the V3 and V4 region of the 16S rRNA gene was amplified with the primer set 515F (5’–GTGCCAGCMGCCGCGGTAA-3’) and 926R (5’– CCGYCAATTYMTTTRAGTTT-3’), [34] to check for the amplification of 16S using the extracted DNA. Illumina DNA libraries and sequencing were performed at Biotika (www.biotika.com.br, Brazil) on a MiSeq platform in a paired-end read run (2 × 250bp) following the manufacturer’s guidelines. Sequencing outputs were the raw sequence data.

### Sequencing Data Processing and Statistical Analyses

The demultiplexed sequences were analyzed with the software package Quantitative Insights Into Microbial Ecology (QIIME 2) version 2019.4 [3]. Sequences were denoised using DADA2 [6] with the following parameters: trim left-f = 19, trim left-r = 18, trunc-len-f = 287, trunc-len-r = 215. Amplicon sequence variants (ASVs) with sequences less than 10 occurrences were removed. The taxonomy was assigned to the representative sequences of ASVs using a Naive Bayes classifier pre-trained on SILVA release 132 clustered at 99% identity. FastTree and MAFFT [17] were used to create a rooted phylogenetic tree which was used in calculating phylogenetic diversity metrics.

Diversity and phylogenetic analyses were performed with the PhyloSeq [25], ggplot2 [45] and vegan [32] packages in R software (Team, 2018). ASVs without at least two sequences occurring in two or more samples and chloroplasts were removed. Alpha-diversity metrics (e.g., observed sequence variants, Chao1 and Shannon diversity) were calculated based on ASV relative abundances for each sample. To determine if there were significant differences between alpha diversities, Analyses of Variance (Kruskal-Wallis) test was performed in R. ASVs were normalized using the R package “*DESeq2*” by varianceStabilizingTransformation [22]. Beta-diversity among sample groups was examined an ordinated weighted Unifrac normalized distance, tested with Permutational Homogeneity of Group Dispersions, and visualized using principal coordinate analysis (PCoA, package Phyloseq). Relative abundance of taxonomic indicators was identified by the IndicSpecies [5]. The analysis was conducted on ASV counts excluding ASVS < 20 reads. Predicted microbial functional groups were identified by the Functional Annotation of Prokaryotic Taxa (FAPROTAX) database [23]. Statistical tests were considered significant at *p* < 0.05. The sequencing reads generated for this study can be found in the National Centre for Biotechnology Information (NCBI) database under the BioProject PRJNA638744.

## RESULTS

### Alpha and Beta Diversity Estimates at Rio Grande Rise

A total of 409,346 sequences was retrieved from eleven samples and clustered into 446 amplicon sequence variants (ASVs) (0.03 cut-off). The number of sequences ranged from 10,347 to 94,103 and the ASVs ranged from 10 to 73 per sample (Supplementary Table 1). In general, mean richness (366 ±159, standard error, s.e.) and diversity (5.38 ± 0.45, s.e.) were higher in calcarenite samples when compared to those from Fe-Mn crust biofilm, Fe-Mn crust and Fe-Mn encrusted coral skeleton from the RGR (Fig. 2 and Supplementary Table 1). Richness (163±140, s.e.) and diversity (3.16±1.27, s.e.) were lowest at Fe-Mn crust community compared to the other substrates with greater variability between samples (see standard error, Fig. 2 and Supplementary Table 1). Although diversity was higher on Fe-Mn coral samples (3.93±0.79, s.e.) compared to the crust biofilm (3.3±0.24, s.e.), the crust biofilm (230.8±55.56, s.e.) community showed a higher richness than the coral (105.5±59.50, s.e.) (Fig. 2 and Supplementary Table 1). Fe-Mn substrates additionally showed the variation in richness and diversity between samples, as evaluated with standard error. Alpha diversity indexes were not statistically different between Fe-Mn substrates (Supplementary Table 1).

**Fig. 2.**
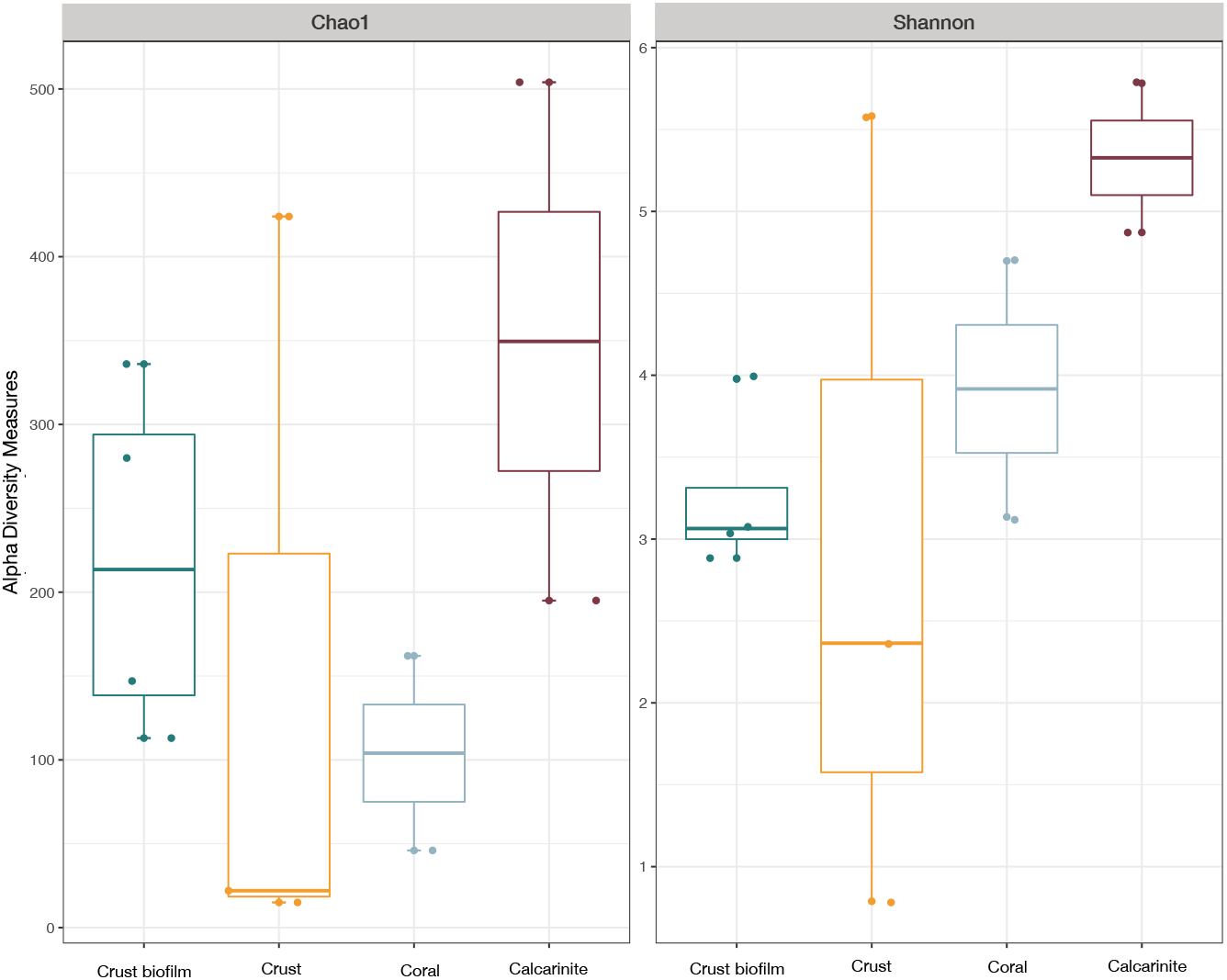
Alpha diversity medians (number of ASVs, Chao1 and Shannon indexes) of microbial communities in the Fe-Mn crust biofilm-like, crusts, encrusted coral skeletons and calcarenites on the RGR

Beta diversity among the samples and substrates was tested using the weighted Unifrac distances and ordered by PCoA. In the PCoA plot, samples were clustered by substrates, expected DY94_56_01 and DY94_46_03 (Fig. 3). The PCoA analysis captured 60.5% of the total variation of the prokaryotic community composition in the investigated samples. Samples were also clustered by depths (Supplementary Fig. 2). The analysis of groups dispersions showed a significant heterogeneity of dispersion among stations (ANOVA, P < 0.05) and depths (ANOVA, P < 0.05).

**Fig. 3.**
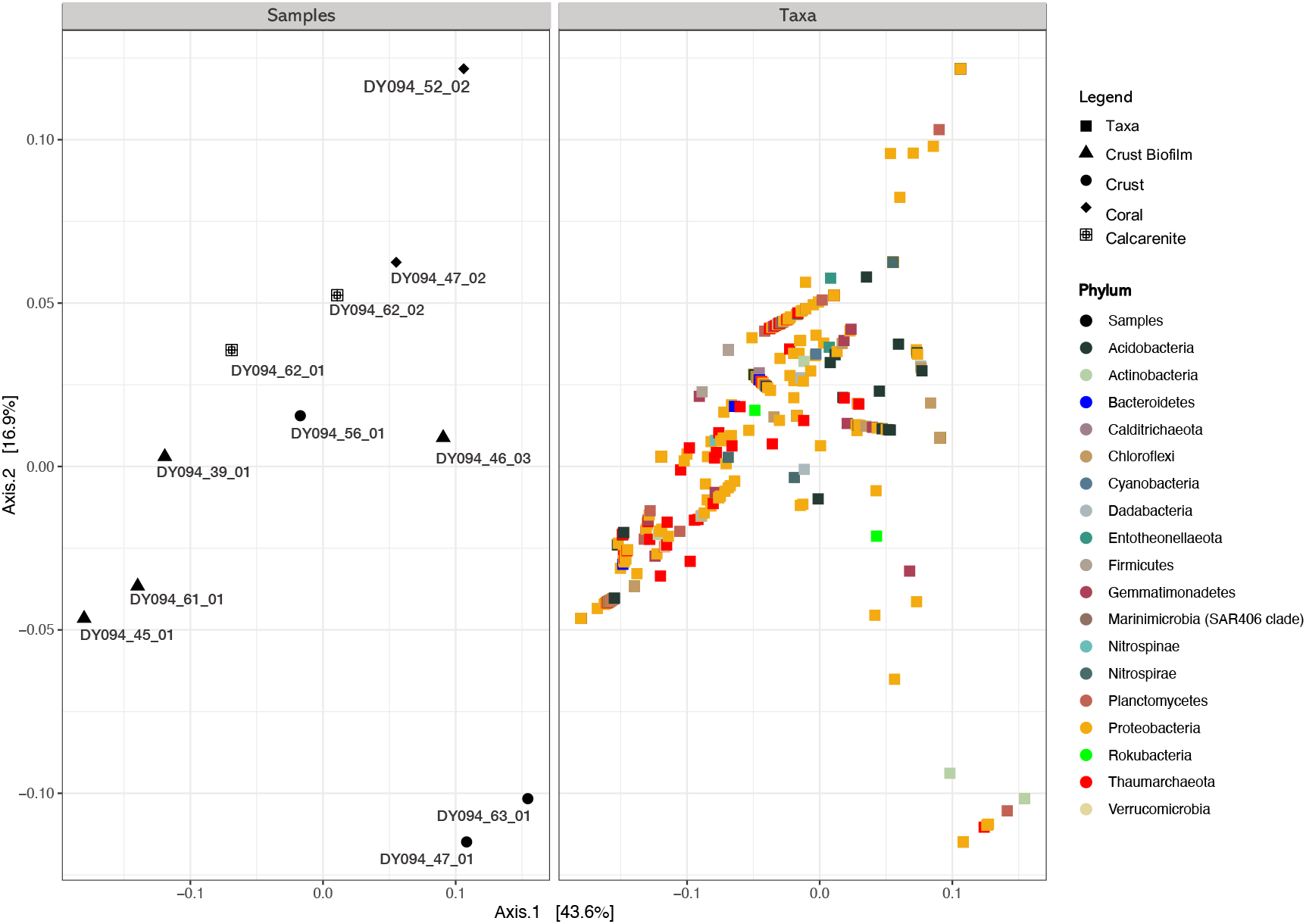
Principal coordinate analysis (PCoA) of the microbial community of Fe-Mn crust biofilm, crusts, encrusted coral skeletons and calcarenites from the RGR. The results were based on the amplicon sequencing data of the 16SrRNA genes using weighted Unifrac distance. The taxa correlating with the community differences (at phylum level) are also shown in the plot on the right (Taxa)

### Microbial Community Composition at Rio Grande Rise

Bacterial and Archaeal composition varied in abundance or occurrence among substrates and samples. For example, the microbial groups Proteobacteria (classes Gammaproteobacteria and Alphaproteobacteria), Thaumarchaeota (Nitrosopumilales) and Planctomycetes (classes Phycisphaerae and Planctomycetacia) were abundant in all substrates, except the DY094_63_01_MB, DY094_47_01_MB, DY094_52_02_MB, DY094_39_01_A and DY094_46_03_MB samples (Fig. 4). Proteobacteria was the most abundant phyla, comprising about 39% of the total taxonomic composition of each sample (Fig. 5). Within Proteobacteria, Gammaproteobacteria was the second most abundant class (22%), followed by Alphaproteobacteria (12%). Nitrososphaeria was the most abundant class (23%), especially among the crust biofilm samples (Fig. 4 and Fig. 5). ASVs affiliated to Verrucomicrobia, Marinimicrobia (SAR406 clade), Calditrichaeota, Cyanobacteria and Firmicutes represented less than 1% of all sequences (Fig. 4A).

**Fig. 4.**
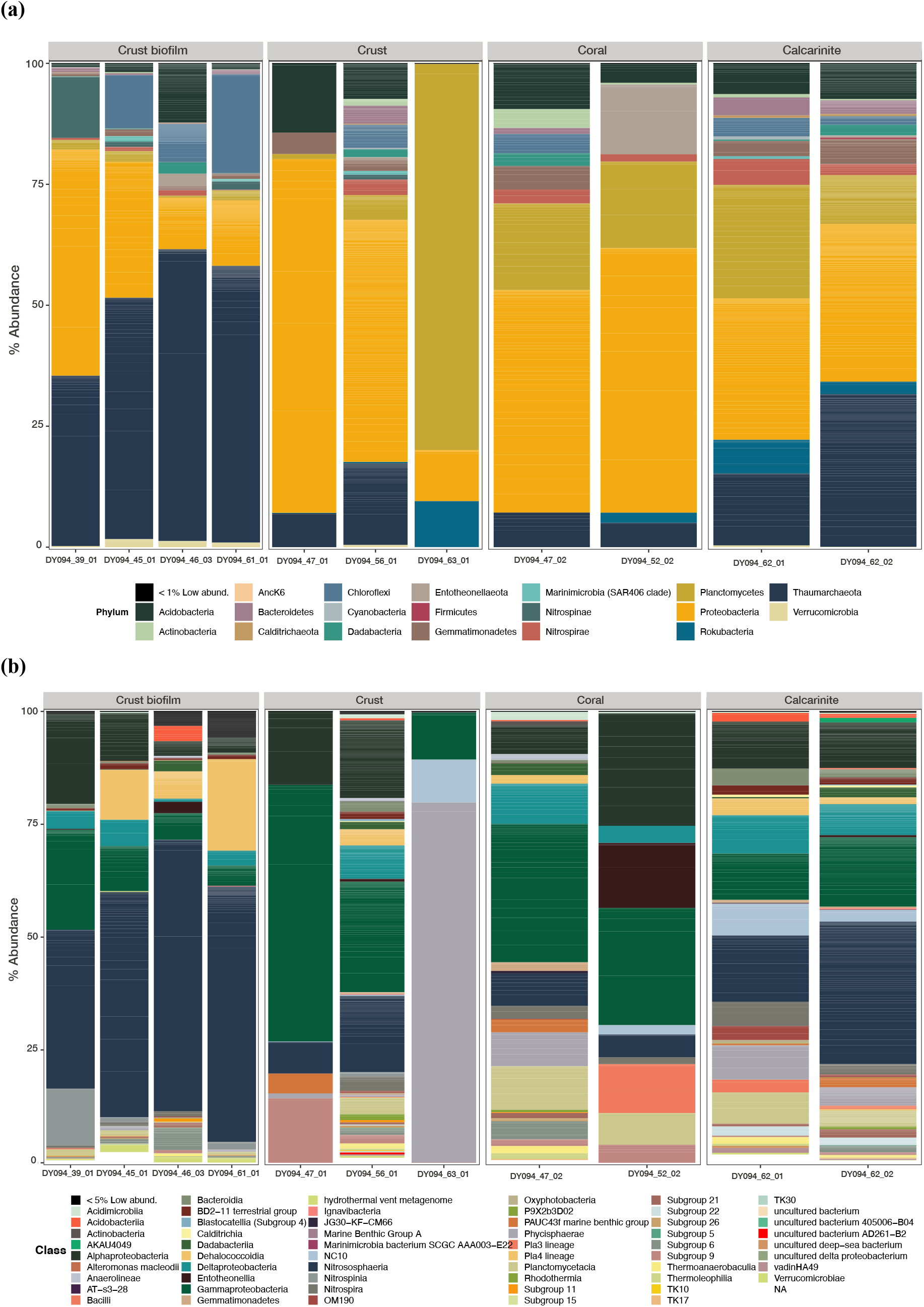
The relative abundances of bacterial and archaeal taxonomic composition for (**a**) phylum and (**b**) class in the Fe-Mn crust biofilm, crusts, encrusted coral skeletons and calcarenites from the RGR. Only phyla and classes with more than 0.1% of abundance are represented. Phyla with relative abundances below 1% and classes with relative abundances below < 5% were grouped together into low abundance groups. Gray boxes at the top indicate sample substrates, i.e. crust biofilm, crust, encrusted coral skeleton and calcarenite

**Fig. 5.**
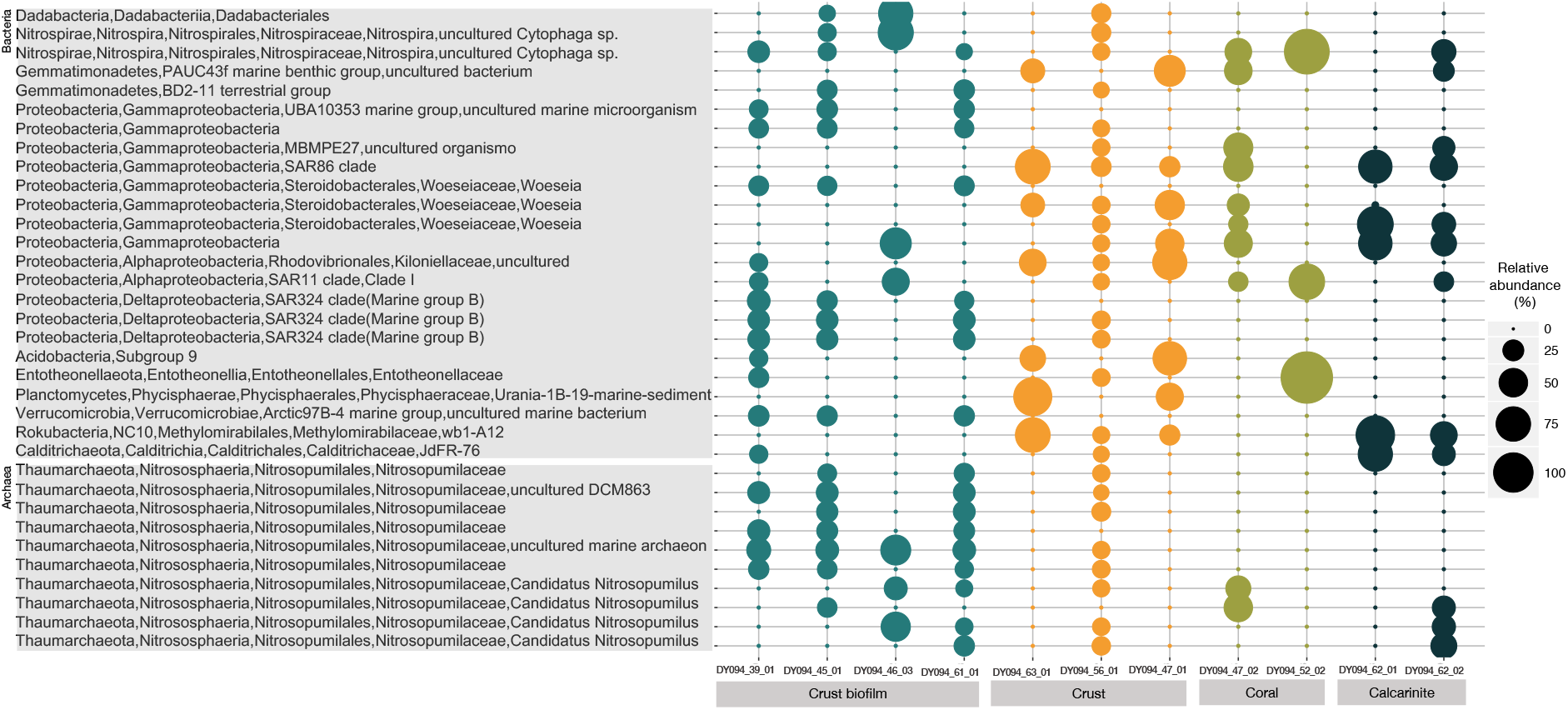
Classification of the most abundant ASVs for Archaea and Bacteria (> 0.1% relative abundance) in the Fe-Mn crust biofilm, crusts, encrusted coral skeletons and calcarenites. The size of circles is related to the relative abundance of each ASV. ASVs are organized by phylum. Gray boxes at the bottom indicate sample substrates, i.e. crust biofilm, crust, encrusted coral skeleton and calcarenite

Thaumarchaeota (55%) was prevalent in crust biofilm samples (Fig. 5), followed by the phyla Proteobacteria (17%), Chloroflexi (12%) and Acidobacteria (6%) (Fig. 4A). ASVs affiliated to Bacteroidetes, Marinimicrobia (SAR406 clade), Actinobacteria, Calditrichaeota, Cyanobacteria and Rokubacteria represented less than 1% of all sequences in crust biofilm samples (Fig. 4A). At the class level in crust biofilm, Nitrososphaeria (60-51%, Nitrosopumilales), Dehalococcoidia (20-7%, order SAR202 clade), Gammaproteobacteria (10-5%, orders Cellvibrionales, Nitrosococcales, UBA10353 marine group and Oceanospirillales), Alphaproteobacteria (10-3%, orders Rhodospirillales, Rhodovibrionales and Sar11) and Deltaproteobacteria (6-1%, orders SAR324 clade and NB1-j) were dominant, except in the DY094_39_01 sample. The bacterial class Dehalococcoidia was not identified in DY094_39_01; the prevalent classes were Gammaproteobacteria (22%), Alphaproteobacteria (20%), Nitrospinia (13%) and Deltaproteobacteria (4%).

The prevalent ASVs in the Fe-Mn crusts were Proteobacteria vAcidobacteria (14-7%) and Thaumarchaeota (17-7%) and Planctomycetes (5-1%), except in the DY094_63_01 sample. The bacterial phyla Planctomycetes dominated (>70%) in DY094_63_01 crust, followed by Proteobacteria (11%) and Rokubacteria (10%) (Fig. 4 and Fig. 5). Representatives of the phyla Chloroflexi, Bacteroidetes, Nitrospirae, Dadabacteria, Actinobacteria and Nitrospinae were detected in crusts only in the DY094_56_01 sample. At the class level, Gammaproteobacteria (57-24% orders Acidiferrobacterales, Alteromonadales, MBMPE27, pItb-vmat-80 and Steroidobacterales), Alphaproteobacteria (17-16%, orders Kordiimonadaceae, Rhizobiales and Rhodovibrionales), Subgroup 9 (14-2%) and Nitrososphaeria (17-7%, Nitrosopumilales) were abundant in DY094_47_01 and DY094_56_01 crusts. Otherwise, representatives of Phycisphaerae (80%, order Phycisphaerales), Gammaproteobacteria (10%, Sar86 clade) and NC10 (9%, order Methylomirabilales) were more abundant in the DY094_63_01 crust (Fig. 4A). ASV related to a Colwelliaceae family were detected in the DY094_47_01 and DY094_63_01 crust samples.

In Fe-Mn encrusted coral skeletons, microbial communities were dominated by ASVs affiliated to Proteobacteria (54-46%), Planctomycetes (17-17%), Thaumarchaeota (7-5%), Acidobacteria (9-4%) and Nitrospinae (3-1%) (Fig. 4A). The prevalent class in coral samples were related to Gammaproteobacteria (27% orders Betaproteobacteriales, MBMPE27, BD7-8 and Steroidobacterales), Alphaproteobacteria (20%, orders Kordiimonadaceae, Rhizobiales, Rhodovibrionales and Sneathiellaceae), Planctomycetacia (7%, order Pirellulales), Nitrososphaeria (5%, Nitrosopumilales) and Deltaproteobacteria (5%, orders Myxococcales and NB1-j), (Fig. 4A). The classes Entotheonellia (14%, order Entotheonellales) and NC10 (2%, order Methylomirabilales) were only detected in the DY094_52_02_MB sample (Fig.5). Representatives of the classes PAUC43f marine benthic group (5%), Dehalococcoidia (4%), Acidimicrobiia (4%), Dadabacteriia (3%), Bacteroidia (1%) and Actinobacteria (0.1%) were detected only in coral from DY094_47_02_MB (Fig. 4B). ASVs related to a Geodermatophilaceae family were detected only in the DY094_47_02 sample.

Calcarenite samples were dominated by ASVs affiliated to Proteobacteria (30%), Thaumarchaeota (20%) Planctomycetes (10%), Acidobacteria (7%), Rokubacteria (5%), Nitrospirae (4%) and Gemmatimonadetes (4%). ASVs associated with Actinobacteria, Calditrichaeota, Nitrospinae, Verrucomicrobia represented less than 1% of all sequences in calcarinete samples (Fig. 4A). At the class level, the prevalent groups were Nitrososphaeria (21%, Nitrosopumilales), Gammaproteobacteria (12%, orders Betaproteobacteriales, Gammaproteobacteria Incertae Sedis, MBMPE27, BD7-8 and Steroidobacterales), Alphaproteobacteria (10%, order Rhodovibrionales), Deltaproteobacteria (8%, orders Myxococcales and NB1-j), Phycisphaerae (6%), Planctomycetacia (6%, order Pirellulales) and NC10 (6%, order Methylomirabilales) (Fig. 4B and Fig. 5). Representatives of Cyanobacteria (class Oxyphotobacteria) and Marinimicrobia (clade SAR406) were detected in the calcarenite sample only from DY094_62_01_MB (Fig. 4B).

Microbial community composition among different substrates was further investigated by means of the Indicator Species analysis. We compared the relative abundance and relative frequency of each ASV to identify those specifically associated with only one substrate (unique) and those whose niche breadth encompasses several substrates (shared). The results were visualized as a table showing the distribution of unique and shared ASV among different substrates (Supplementary Table 2). Calcarinite harbored the highest set of unique ASVs (n = 24), mainly belong to the oligotypes Nitrosopumilales (n = 6, Thaumarchaeota), class of uncultured Alphaproteobacteria (n = 5), class Gammaproteobacteria (n = 3, families EPR3968-O8a-Bc78v and Nitrosococcaceae), class Phycisphaerae (n = 2, family Phycisphaeraceae), class Nitrospira (n = 2, family Nitrospiraceae) and class Gemmatimonadetes (n = 2, family Gemmatimonadaceae and class PAUC43f marine benthic group) (Supplementary Table 2).

However, crusts, crust biofilm and encrusted coral skeletons harbored fewer unique ASVs each (n = 1, n = 1 and n = 2, respectively). Those ASVs that are unique to crust and crust biofilms were ASV1055 and ASV0706, belonging to the class Gammaproteobacteria and Nitrosopumilales (Thaumarchaeota), respectively. The unique ASVs of encrusted coral skeletons are ASV0714 and ASV1608, from the family Pirellulaceae (phylum Planctomycetes) and the order Rhizobiales (class Alpaproteobacteria). Crust and Calcarenite share only one ASV that was related to the order Methylomirabilales (class NC10, ASV0807) (Supplementary Table 2).

### Predicted Function Variation among the Microbial Communities

We assigned 561 out of 1875 microbial ASVs (29.92%) to at least one microbial functional group using the FAPROTAX database (Supplementary Table 3). Aerobic ammonia oxidation, aerobic nitrite oxidation and nitrification were functions predicted in all substrates with differences in relative abundance between samples (Fig. 6). Main functions associated with crust biofilm microbial communities were dark sulfide oxidation, dark oxidation of sulfur compounds, nitrate respiration, nitrate reduction, nitrogen respiration and aerobic chemoheterotrophy. The predicted functions in crusts and calcarenite samples were fermentation and aerobic chemoheterotrophy. Manganese oxidation was predicted only in the DY094_47_02 coral sample (Fig. 6).

**Fig. 6.**
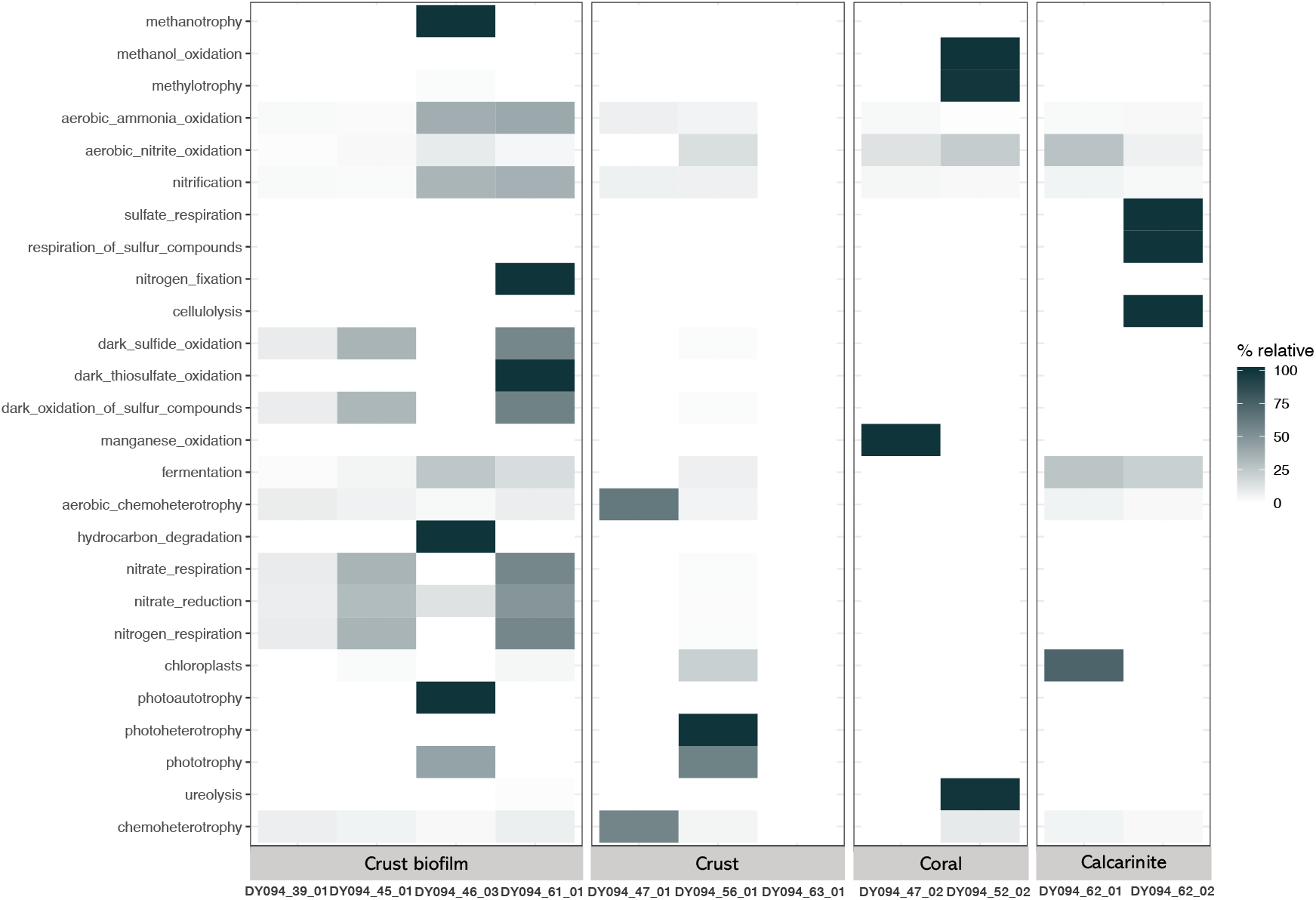
Mean of relative abundances of microbial functional groups in Fe-Mn crust biofilm, crusts, encrusted coral skeletons and calcarenites. Relative abundances are depicted in terms of color intensity from white (0) to dark green (100)

## DISCUSSION

Our results showed that bacterial and archaeal community structure in Fe-Mn crusts are less diverse in comparison with other substrates: crust biofilm, encrusted coral skeleton and calcarenite. We also observed differences in microbial community structure between the sampling stations and depth, as previously reported by Kato et al. (2018) for the Fe–Mn crusts from the Pacific Ocean. Three factors might be shaping the community structure and composition variance on the Fe–Mn substrates from the RGR. The first is related to the water depth and location, because the concentrations of elements in the crusts change depending on the water depth [1, 41]. The second might be related to the geochemical differences in the composition of the studied substrates, which can promote variations in the lifestyles of solid-surface attached microbes [16]. The last factor is probably associated with the local circulation, as previously described for the Fe-Mn deposits from the Pacific Ocean and for the RGR [1, 16, 24, 31]. Further analyses are needed to clarify whether and how the physical oceanography and geochemical composition of Fe-Mn substrates in the RGR are associated with the differences observed in microbial communities.

Microbial assemblages in samples from the RGR showed the dominance of the classes Gammaproteobacteria, Alphaproteobacteria and Deltaproteobacteria, as reported for Pacific Fe-Mn crusts [21, 38, 42, 46]. However, at higher taxonomic resolution, we detected differences in the microbial community composition between samples from the RGR and those from the deep Pacific Ocean. Samples from the RGR showed higher abundance of the orders Steroidobacterales (family Woeseiaceae), uncultured MBMPE27, Methylomirabilales and Nitrospirales, when compared to Fe-Mn crusts from other regions [16, 24].

Previous studies have reported that dissolved ammonia in seawater is an important nutrient in the Fe-Mn crusts from the Pacific Ocean [15, 16, 30, 31]. Based on the recovered ASVs and the functional predictions performed in this study, nitrification (i.e., ammonia oxidation and nitrite oxidation) and carbon fixation might be the main processes occurring in the Fe-Mn substrates from the RGR. We detected a high proportion of chemolithoautotrophic ammonia-oxidizing Archaea (Nitrososphaeria class) in the Fe-Mn substrates, especially in our Fe-Mn crust endolithic biofilm samples, as well as chemolithoautotrophic nitrate-oxidizing Bacteria within Nitrospirae and Nitrospinae. High proportions of these archaeal ammonia oxidizers (class Nitrososphaeria) and bacterial nitrate, urea and cyanate oxidizers (Nitrospirae and Nitrospinae) have also been reported in abyssal Fe– Mn deposits in the South Pacific Gyre [15], in the Clarion and Clipperton Zone [30, 31, 38] and in the Peru basin [26].

Shiraishi et al. (2016) suggested that Nitrososphaeria might be involved in manganese oxidation in Fe-Mn nodules, as they have multi-copper oxidase, a gene that is utilized by most known manganese oxidizers. Also, previous study proposed that Mn-reducing bacteria, such as species within *Shewanella* and *Colwellia*, are the major contributing microorganisms in the dissolution of Mn in Fe–Mn crusts [2]. Although we have not detected ASVs related to *Shewanella,* we identified ASVs in our results associated with acetate-oxidizing manganese reducers (Colwelliaceae) and manganese oxidizers (Geodermatophilaceae). These manganese oxidizers and reducers were in low abundance and detected only in Fe-Mn crusts and encrusted coral skeletons from stations 47 and 63. Further, we detected a low abundance of ASVs in the Fe-Mn crust biofilm associated with iron reducers from the Magnetospiraceae family. Magnetospiraceae and Colwelliaceae families have been reported previously across the Pacific Nodule Provinces and the Peru Basin [26, 38]. Other possible mechanisms for Mn release on the Fe-Mn substrate surface are acidification through the production of nitrite by ammonia oxidizers and organic acid production by fermenting microorganisms [16]. Indeed, we detected ASVs related to ammonia oxidizers and fermenting microorganisms, including members of the orders Bacillales, Pseudomonadales and Azospirillales.

Biofilms with filamentous microorganisms associated with micro-stromatolitic growth bands have been reported in the Fe-Mn nodules and crusts surfaces from Pacific Ocean [4, 13, 40, 42]. Recently, Thaumarchaeota affiliated metagenome-assembled genome (MAG) was recovered from Fe-Mn crust [15] and proteins associated with biofilm formation and surface adhesion capabilities were detected [15]. Kato et al (2019) suggested that these biofilms probably contribute to the Fe-Mn crust formation by adsorbing Fe and Mn oxide particles from seawater [15]. Moreover, endolithic microorganisms within coral skeletons have been previously described [47], but those associated with Fe-Mn substrates are poorly studied. We found within the microbial community of our Fe-Mn encrusted coral skeleton an ASV belonging to the order Phycisphaerales (phylum Planctomycetes), which was previously associated with tropical stony corals found mainly in cold waters [18]. Gammaproteobacteria are common members of coral microbiomes [18]. Our main contributing gammaproteobacterial family were related to the Woeseiaceae (order Steroidobacterales), in contrast to other studies of deep-sea corals [18, 47]. Members of the Woeseiaceae family were abundant in all Fe-Mn substrates in this study. The Woeseiaceae family is cosmopolitan in deep-sea sediments [29] and was also reported across an Fe-Mn nodule field in the Peru basin [26]. The fact isolated Woeseiaceae members show a chemoorganoheterotrophic lifestyle [8], suggested a role in organic carbon remineralization in the deep-sea [11].

In our Fe-Mn substrates, we also detected higher abundance of ASVs belonging to methane oxidation, i. e. Methylomirabilales and SAR324. Members of the order Methylomirabilales, within phylum NC10, were abundant in Fe-Mn crusts and calcarenite, and are capable of nitrite-dependent anaerobic methane oxidation (Versantvoort et al., 2018). SAR324 of the Deltaproteobacteria was abundant in our crust biofilm and crusts and is known to have wide metabolic flexibility, including carbon fixation through Rubisco and an ability to oxidize alkanes, methane and/or sulfur to generate energy [20, 36].

In addition to carbon, the results indicate that nitrogen metabolism is important in Fe-Mn substrates from the RGR. This is probably because the RGR is in the oligotrophic South Atlantic Gyre, with a low concentration of organic carbon and high concentrations of nitrate, phosphate and Fe-Mn substrates [27]. Organic carbon compounds in the deep-sea might be recalcitrant for microbial life [14], whereas Mn oxides are known to abiotically decompose recalcitrant organic carbon (i.e., humic substances) to simple carbon compounds, such as pyruvate, acetaldehyde, and formaldehyde [39]. These simple carbon compounds can be used as carbon sources by deep-sea microorganisms, as members of family Woeseiaceae and classes Methylomirabilales and SAR324 [39].

Finally, this study is the first to report microbial diversity in Fe-Mn substrates from the deep Atlantic Ocean. Our results reveal that: (1) the microbial community compositions among Fe–Mn substrates are different according to the water depths and sampling location, (2) populations associated with nitrogen and carbon metabolisms are likely important contributors for the ecological process occurring in Fe-Mn substrates, (3) the microbial community composition detected on the RGR are not similar to those described in Fe-Mn crusts from the Pacific Ocean, except for a higher proportion of ammonia-oxidizers, which suggest the commonality predominance of this group in Fe-Mn substrates from both the Pacific and Atlantic oceans, and (4) a similar microbial community composition was detected in an oligotrophic Fe-Mn nodules field from Peru basin, i.e. Woeseiaceae, Methylomirabilales, Rhodovibrionales, BD2-11 terrestrial group and SAR324, Subgroup9. Thus, further monitoring of the benthic-pelagic coupling microbiome, such as metagenomics and metatranscriptome, and measurement of the oceanographic conditions in the RGR are needed to better elucidate the ecological processes involved with the formation of Fe-Mn substrates.

## Acknowledgments

We thank the captain and the crew of the Royal Research Ship Discovery cruise DY094 for their data and sampling support, as well as LECOM's research team and Rosa C. Gamba for their scientific support.

## SUPPLEMENTARY MATERIAL

**Supplementary Figure 1.**
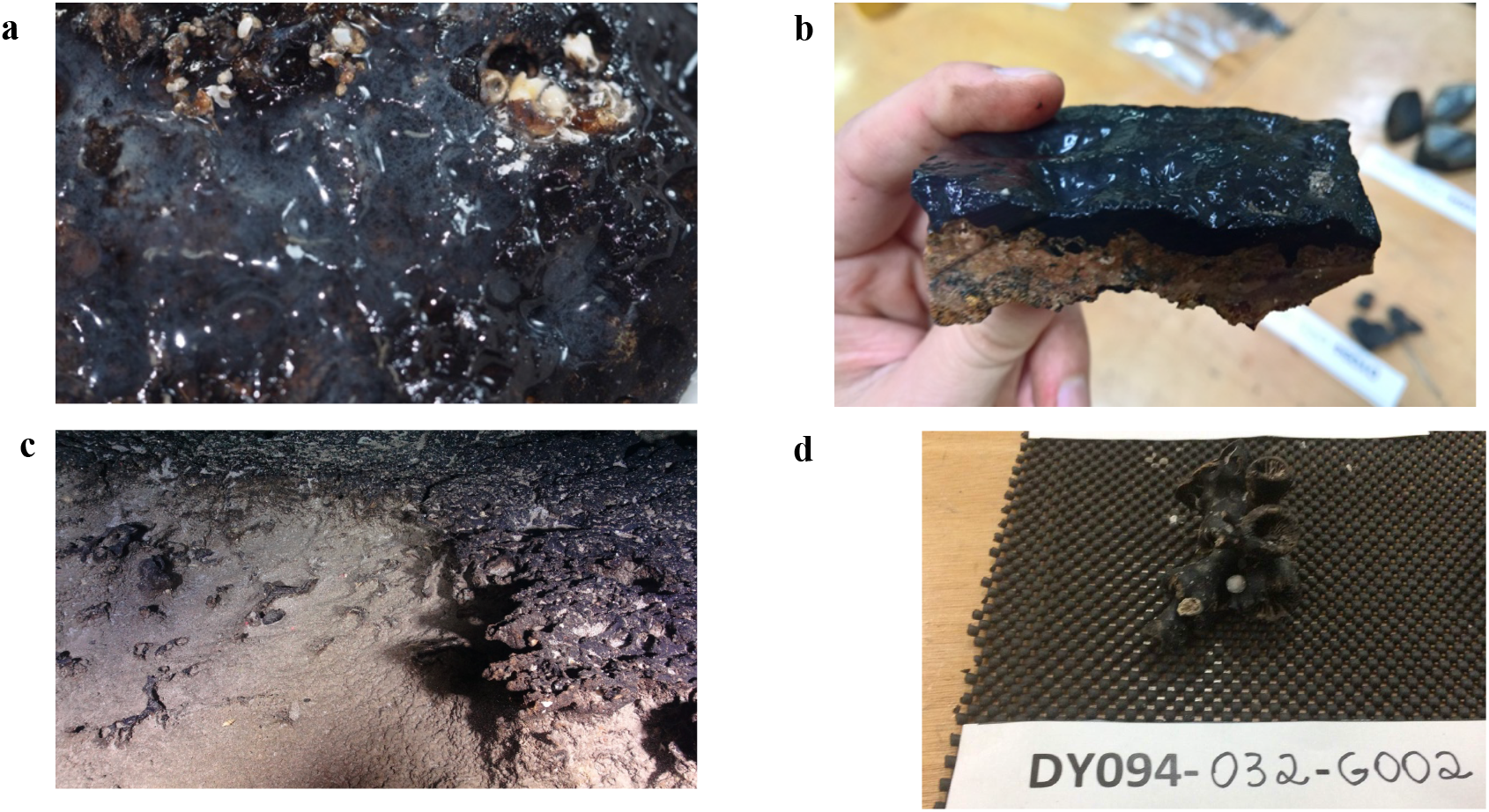
Examples of (**a**) Fe-Mn crust biofilm on Fe-Mn crust, (**b**) Fe-Mn crust, (**c**) calcarenite and (**d**) Fe-Mn encrusted coral skeletons collected during the scientific expedition on the RRS Discovery on the RGR

---

**Supplementary Table 1.**
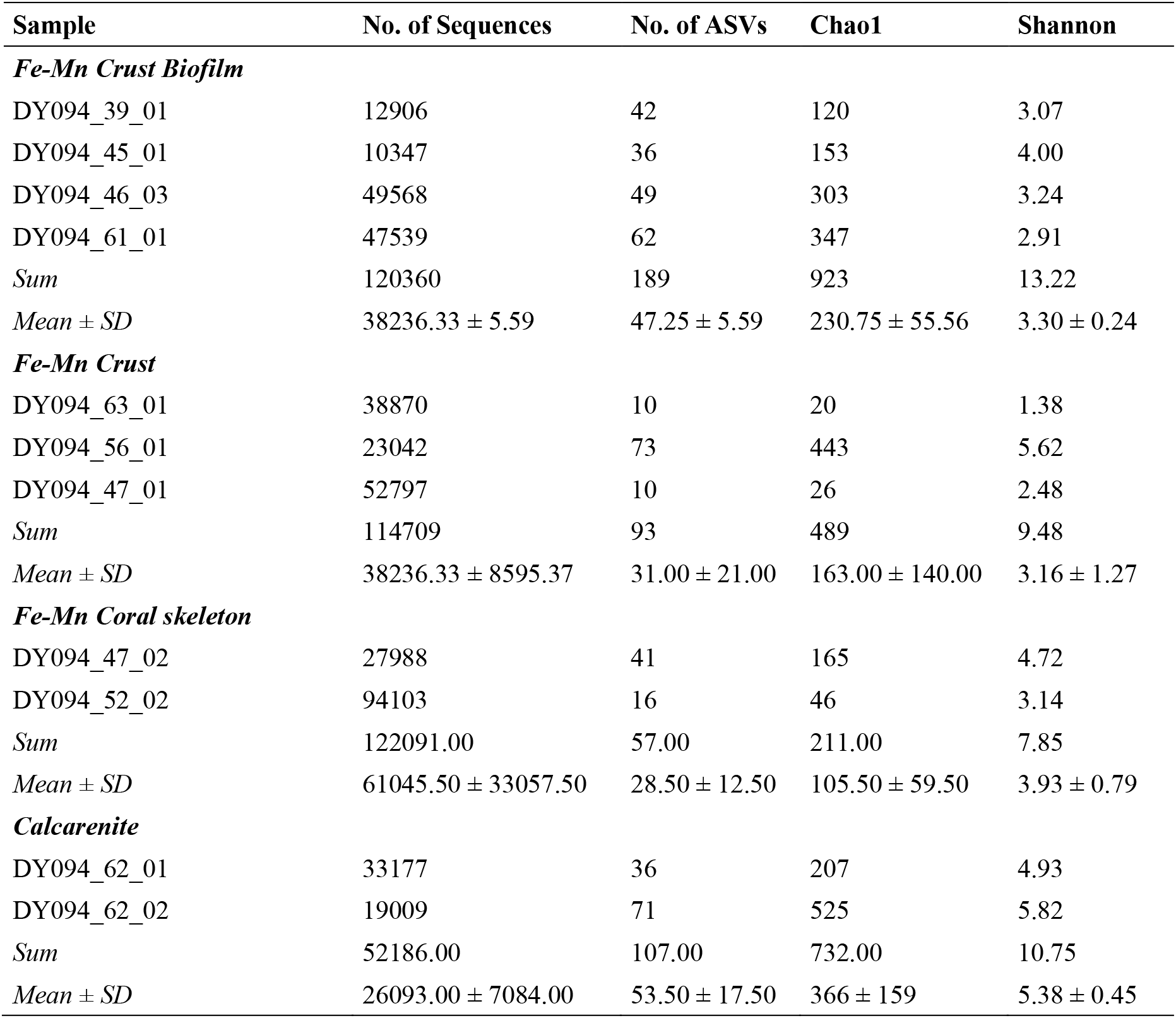
Sum, mean, standard error (S. error) for number of sequences, number of amplicon sequence variants (ASVs), Chao1 and Shannon indexes for Fe-Mn crust, crust biofilm, encrusted coral skeleton and calcarenite from the RGR.

**Supplementary Table 2.**
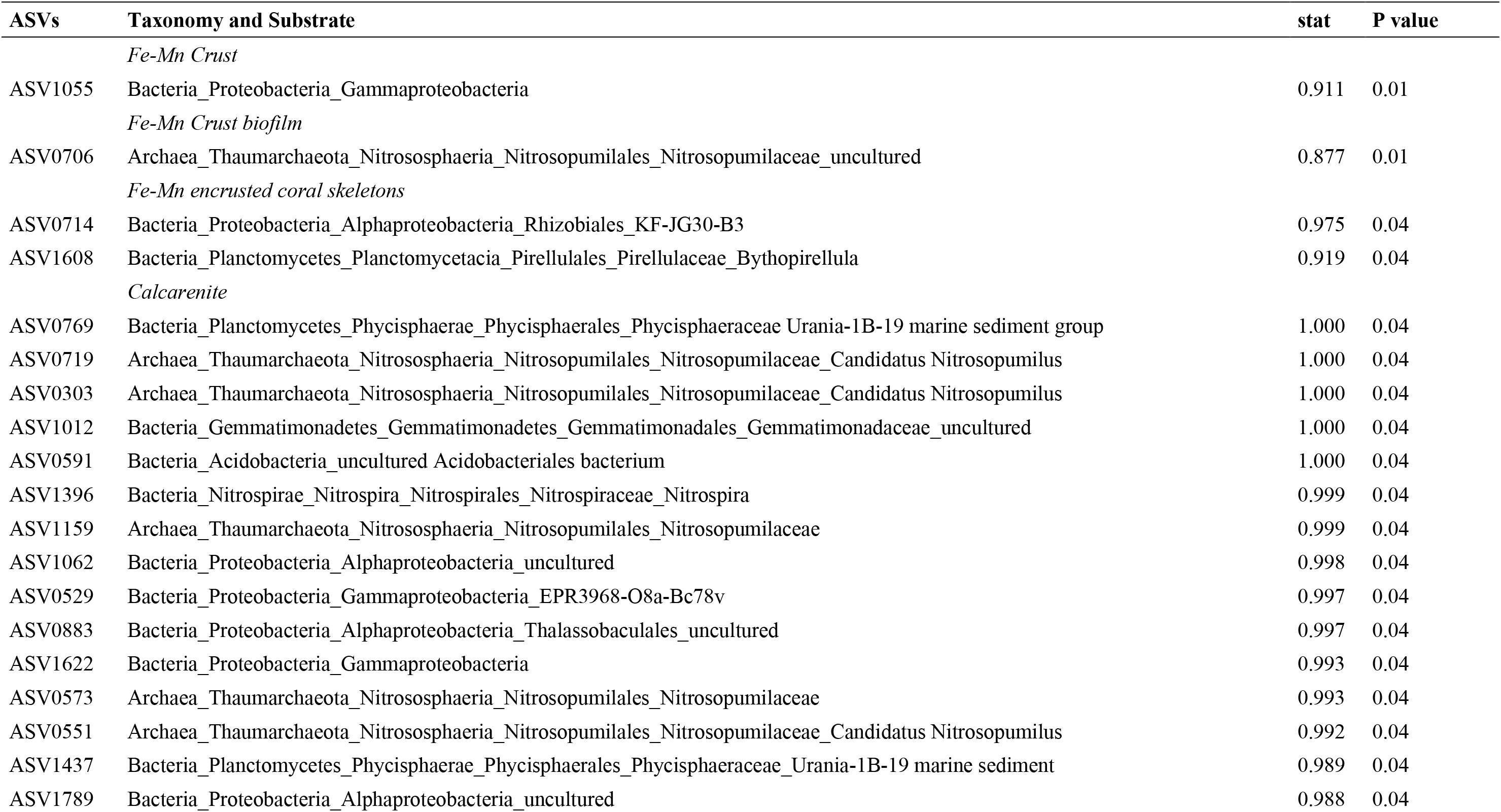
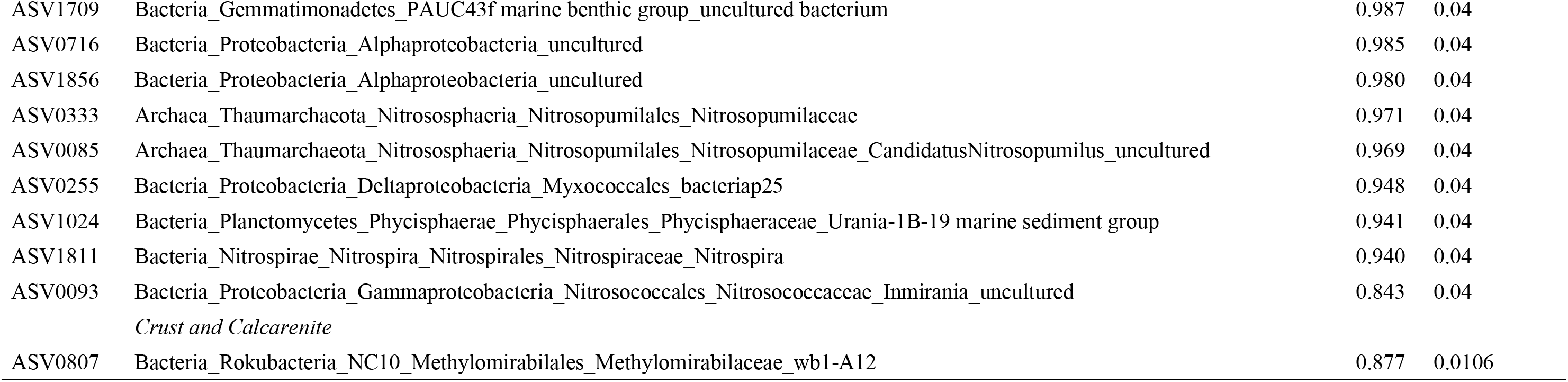
Index Species result with significant amplicon sequence variants (ASVs) and taxonomy organized by substrate.

**Supplementary Table 3.** Microbial functional groups predicted in the Fe-Mn crust biofilm, crust, encrusted coral skeleton and calcarenite from the RGR, and the number of bacterial and archaeal amplicon sequencing variants (ASVs) identified for each functional group.

| Microbial Function Predicted | ASVs assigned |
| --- | --- |
| chemoheterotrophy | 119 |
| aerobic_chemoheterotrophy | 113 |
| nitrification | 124 |
| aerobic_ammonia_oxidation | 81 |
| aerobic_nitrite_oxidation | 43 |
| predatory_or_exoparasitic | 10 |
| intracellular_parasites | 10 |
| fermentation | 9 |
| animal_parasites_or_symbionts | 9 |
| human_pathogens_all | 8 |
| nitrate_reduction | 3 |
| dark_oxidation_of_sulfur_compounds | 3 |
| nitrogen_fixation | 2 |
| dark_sulfide_oxidation | 2 |
| nitrogen_respiration | 2 |
| nitrate_respiration | 2 |
| phototrophy | 2 |
| ureolysis | 2 |
| chloroplasts | 2 |
| methylotrophy | 2 |
| sulfate_respiration | 1 |
| respiration_of_sulfur_compounds | 1 |
| hydrocarbon_degradation | 1 |
| manganese_oxidation | 1 |
| dark_thiosulfate_oxidation | 1 |
| cellulolysis | 1 |
| fish_parasites | 1 |
| methanotrophy | 1 |
| methanol_oxidation | 1 |
| cyanobacteria | 1 |
| oxygenic_photoautotrophy | 1 |
| photoautotrophy | 1 |
| photoheterotrophy | 1 |

